# Predicting Dog Phenotypes from Genotypes

**DOI:** 10.1101/2022.04.13.488108

**Authors:** Emily R. Bartusiak, Míriam Barrabés, Aigerim Rymbekova, Julia Gimbernat-Mayol, Cayetana López, Lorenzo Barberis, Daniel Mas Montserrat, Xavier Giró-i-Nieto, Alexander G. Ioannidis

## Abstract

We analyze dog *genotypes* (*i*.*e*., positions of dog DNA sequences that often vary between different dogs) in order to predict the corresponding *phenotypes* (*i*.*e*., unique observed characteristics). More specifically, given chromosome data from a dog, we aim to predict the breed, height, and weight. We explore a variety of linear and non-linear classification and regression techniques to accomplish these three tasks. We also investigate the use of a neural network (both in linear and non-linear modes) for breed classification and compare the performance to traditional statistical methods. We show that linear methods generally outperform or match the performance of non-linear methods for breed classification. However, we show that the reverse is true for height and weight regression. Finally, we evaluate the results of all of these methods based on the number of input features used in the analysis. We conduct experiments using different fractions of the full genomic sequences, resulting in input sequences ranging from 20 SNPs to ∼200k SNPs. In doing so, we explore the impact of using a very limited number of SNPs for prediction. Our experiments demonstrate that these phenotypes in dogs can be predicted with as few as 0.5% of randomly selected SNPs (*i*.*e*., 992 SNPs) and that dog breeds can be classified with 50% balanced accuracy with as few as 0.02% SNPs (*i*.*e*., 40 SNPs).

## I. INTRODUCTION

An organism’s DNA sequence contributes to the way it operates, from basic survival functions to unique appearance traits. Most positions in DNA sequences (*i.e*., genomes) do not vary between individuals of the same species. The small percentage of positions that do vary are called *single nucleotide polymorphisms (SNPs)*. SNPs are typically encoded as binary variables, where *zero* denotes a common variant (the variant found in a majority of specimens) and *one* denotes a minority variant. The frequencies of SNP variants at each DNA position and the correlations between nearby variants change for specimens of different animal breeds and plant cultivars. These variances of SNP frequencies and correlations resulted from selective breeding and demographic histories. In this paper, we develop predictive models (*i.e*., statistical and machine learning techniques) to exploit these variances to infer breeds and phenotypes from genomic sequences.

The adoption of such predictive models is useful in a wide range of applications. Predictive models aid in treatment and drug development for domestic animals (*e.g*., dogs, cats, horses) and selection of desired traits during breeding. Similarly, they enable design of better crops with increased nutritional properties that are more resistant to herbicides, insects, and adverse weather. The same set of predictive techniques are being applied to humans for disease prevention, treatment, and pharmaceutical development. Genomic sequences of crops and animals carry fewer privacy restrictions than human data, so working with crop and animal data provides a more flexible environment to explore techniques that can later be applied to human data.

Rapidly decreasing prices of genomic sequencing technologies enables generation of large datasets with a growing coverage of SNPs (*i.e*., high-resolution sequencing). Such datasets are used in Genome-Wide Association Studies (GWAS), which look for correlations between each position of the chromosome and a given phenotype (*i.e*., trait). Such analyses are typically displayed as Manhattan plots, where a negative log*p*−value is shown at each genomic position. GWAS provides a good basis to find causal relationships between positions (*i.e*., SNPs) and traits. By using coefficients obtained in GWAS, Polygenic Risk Scores (PRS) can be constructed to predict a trait given an input sequence.

The phenotypic variation between dog breeds and their population structure, breeding, and migration history has been studied extensively [1]–[5]. While most of the previous work relies on GWAS to explore trait variation and population genetic clustering techniques to explore breed differences, we utilize supervised machine learning to obtain insights into the relationship between genomic sequences and traits. Specifically, we explore a multitude of machine learning techniques for predicting breeds and phenotypes (height and weight) of dogs from their genomic sequences. Similar studies have been recently proposed: [5] explores multiple machine learning techniques for breed and population prediction along the genome; [6] makes use of matrix decomposition to cluster human and dog data; [7], [8] present autoencoders for dimensionality reduction and classification of dog and human data; and [9], [10] explore neural networks for population clustering and prediction.

The main motivation to investigate machine learning (ML) models for dog breed and phenotype prediction is to determine if a specific family of methods (*e.g*., trees, linear models, neural networks) achieves greater success than others with genomic data. ML methods have different inductive biases that contribute to their capabilities in modeling certain data distributions. While there is some intuition for using ML methods on DNA sequences (*e.g*., trees work well with discrete data; linear models capture differences in mean values and additive relationships; kernel methods compare pairs of sequences), there is not a clear consensus on which ML method works best with DNA (specifically SNP sequences). Furthermore, non-linear ML methods for SNP sequence analysis are largely unexplored. Given the success of ML on a wide variety of tasks and the lack of exploration of ML on SNP data, we investigate ML approaches for dog breed and phenotype prediction. We use multiple classifiers and regressors to better understand which techniques work well with SNP sequences. We find that linear models are highly competitive for breed classification. Additionally, we show that a Multi-Layer Pereceptron (MLP) without activation functions (therefore an overparameterized and regularized logistic regression) outperforms non-linear neural networks. On the other hand, non-linear techniques show strong performance for regression tasks. Recent work confirms these trends [5], [10], [11].

Finally, we show how these models perform when the number of input features (*i.e*., SNPs) is highly reduced. The price of genomic sequencing is proportional to how many DNA positions are sequenced, and genotyping array technologies can be developed to obtain a few specific genomic positions at a very low cost. Therefore, showing that predicting breed and (some) phenotypes can be performed with a very low number of SNPs could impact future designs of genotyping arrays and decrease the cost of sequencing dog DNA. This has a clear benefit for both commercial and research applications. In this paper, we demonstrate that accurate phenotype prediction is possible with only 0.5% of the available SNPs.

## II. METHOD

### A. Dataset

Dogs have 38 pairs of non-sex chromosomes [12], [13] with approximately 2.4 billion genetic positions [14]. We analyze a subset of this genetic code in our experiments. Specifically, we use the same genotyping array employed by the company Embark. Our dataset – derived from [3] – consists of 198,473 SNPs for 482 different purebred dogs of 75 breeds. In other words, each data sample is a sequence of 198,473 input features represented as binary values.

Dog breeds can be organized into clades, which are groups of dog breeds that are believed to have a common ancestor. Because our dataset includes so many dog breeds, we visualize the dataset with clade information to understand higher-level relationships of the clades. Then, we can extend our conclusions to the breeds within the clades. Figure 1 shows the distribution of samples in our dataset based on dog clade and illustrates the unbalanced nature of our dataset, with some clades (and thus breeds) containing significantly more samples than others. For example, the Terrier clade includes 112 samples, while the Mediterranean clade includes only 10 samples.

**Fig. 1:**
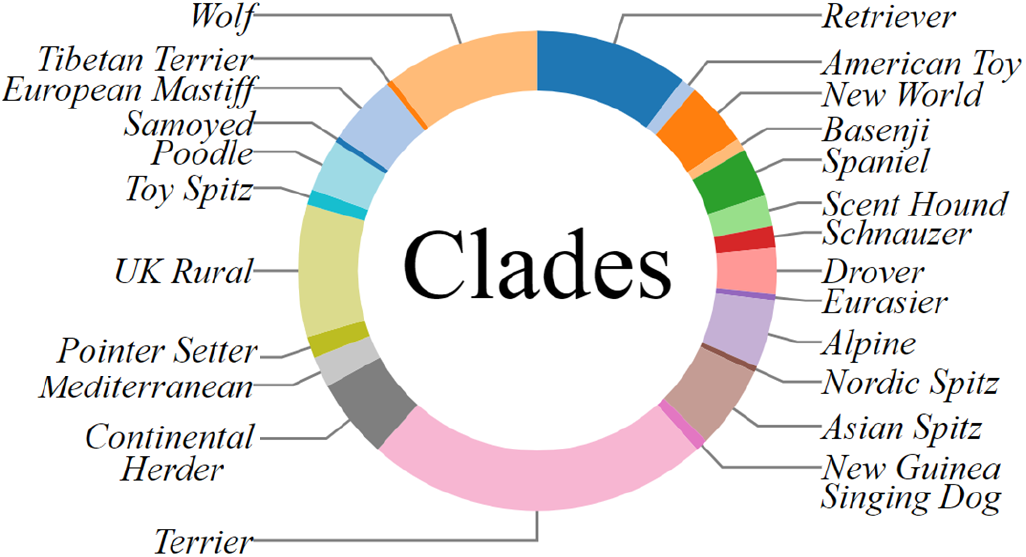
Dog clades in the dataset. Each color represents a different clade, and the size of the colorblock represents the number of dogs in the dataset of that clade.

Figure 2 shows the first two components of a Principal Component Analysis (PCA) [15], [16] and a t-Distributed Stochastic Neighbor Embedding (tSNE) [17] of the SNPs sequences in our dataset. Each data sample is color-coded based on its clade, and some clusters are labeled with breed information. This further shows the unbalanced nature of the dataset in terms of breeds. For example, the Yorkshire Terrier breed occurs 75 times in the dataset, while the Alaskan Husky appears only twice. To ensure that we evaluate breed classification methods on every dog breed, we randomly select one data sample per breed to include in a testing dataset. The training dataset consists of the remaining samples. The resulting training and testing datasets contain 407 and 75 samples, respectively. Only 337 samples in our dataset have height and weight labels (in centimeters and kilograms, respectively), so we use these samples for the height and weight regression tasks. In this case, there are 291 training samples and 46 testing samples. Although it is typical to use data augmentation in machine learning to artificially increase the size of the dataset, it is not common to use data augmentation with SNP sequences. Instead, we employ regularization techniques to help the machine learning methods avoid overfitting.

**Fig. 2:**
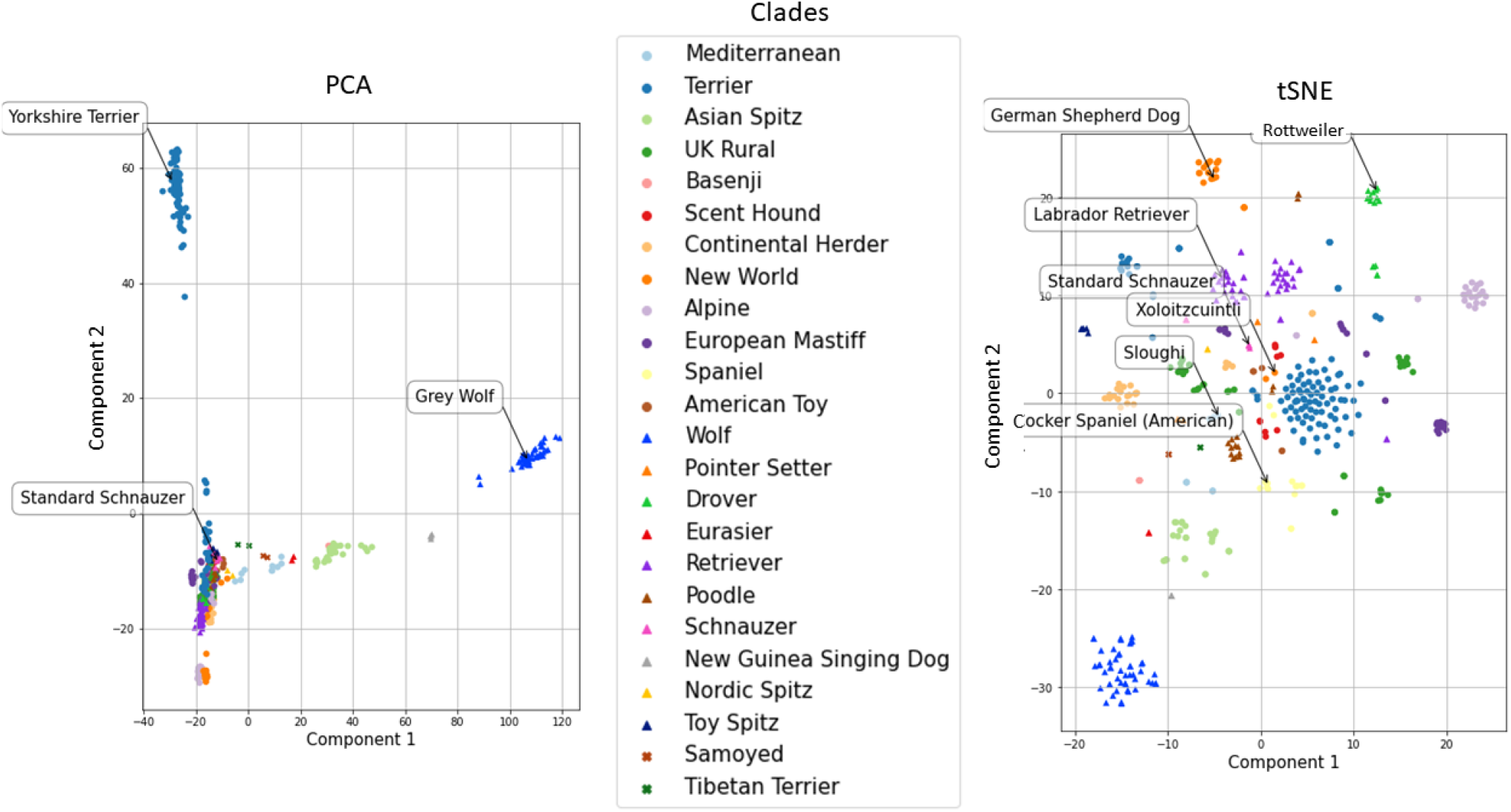
Principal Component Analysis (PCA) and t-Distributed Stochastic Neighbor Embedding (tSNE) of the SNPs sequences in the dataset. Results are color-coded according to dog clade.

Figure 2 also alludes to the complexity of phenotype prediction based on SNPs. Some clades (and thus breeds) are more separable than others. For example, PCA separates Terrier, Asian Spitz, New Guinea Singing Dog, and Wolf clades. Since these clades contain distinctive SNPs sequences, predicting phenotypes of dogs in these clades should be easier than predicting the phenotypes of a Standard Schnauzer. Although a large cluster of overlapping clades occurs in the PCA plot based on two components, analyzing more components could decompse this cluster further. Instead of analyzing each combination of components, we use a tSNE to further separate the data. The tSNE shows a more dispersed visualization of dog clades, enabling a fine-grained analysis of the clusters. We see that the tSNE plot separates the Wolf clade from the other clades, similar to the PCA plot. It also shows distinctive clusters, such as the clusters with German Shepherd Dog and Rottweiler. Thus, these breeds might be easier to classify and predict height and weight for accurately. On the other hand, Xoloitzcuintli dogs might be more difficult to identify because they have so few samples in the dataset without a distinct cluster.

### B. Breed Classifiers and Phenotype Regressors

We explore several multi-class classification methods to identify 75 different dog breeds. We use logistic regression [18]; K-Nearest Neighbors Classifier (KNN) [19], [20]; linear and non-linear Support Vector Machines (SVMs) [21]; Decision Tree Classifier [22], [23]; Random Forest Classifier [24]; AdaBoost [23], [25], [26]; Multi-Layer Percpetrons [23], [27]; Gaussian Naive Bayes; and Gaussian Process Classifier [28]. These methods fall into two major categories: linear and non-linear methods.

To evaluate linear versus non-linear methods more explicitly, we design a Multi-Layer Perceptron (MLP) network that operates in both linear and non-linear modes, depending on whether or not Rectified Linear Unit (ReLU) [29] (*i.e*., a non-linear activation function) is utilized. We also explore the effects of regularization – in the form of batch normalization and dropout – on this task. Through all these experiments, the base architecture of the MLP remains the same: two hidden layers of 1,500 nodes. Only the input layer’s size adapts to the length of the input sequences (*i.e*., from 20 SNPs to 198,473 SNPs). We refer to the base architecture without batch normalization, dropout, or activation functions as *MLP1 (Standard)*. Its over-parameterized architecture provides further regularization and increases the expressive capacity of the network to handle longer input sequences.

We utilize multiple univariate regressors to predict height and weight of the dogs. Again, we explore both linear and non-linear techniques. Specifically, we use Elastic Net (linear regression with L1/LASSO and L2/Ridge regularization); XGBoost; Support Vector Regressors; and K-Nearest Neighbors Regressor. While linear methods could suffice to separate breeds, the relationship between genome and height/weight could be non-linear and require non-linear techniques for accurate phenotype prediction. We conduct a grid search to determine hyperparameters (*e.g*., learning rate, kernel, regularization) used for each method.

## III. RESULTS

In total, we have 198,473 SNPs available per sequence to analyze. However, we wish to investigate how few SNPs are necessary to correctly predict dog phenotypes. Thus, we conduct all our experiments for different fractions of the whole genomic sequence. Let *x* represent a percent of SNPs, where *x* ∈ {0.01%, 0.02%, …, 0.1%, 0.5%, 1%, 5%, 10%, 50%, 100% }. At a minimum, we analyze only 20 SNPs (when *x* = 0.01%), and at a maximum we analyze all 198,473 SNPs available (when *x* = 100%). For every *x*, we conduct 20 experiments with each of the classification and regression methods. Each time a new experiment commences, a new subset of the full SNPs sequences is randomly sampled. To obtain the results shown in Figures 3 and 4, we average the results of the 20 experiments per method and per percentage and calculate the standard deviation of the experimental results. The small standard deviation bars in our figures indicate that there is not much variation in the randomly selected SNPs. A proper selection of SNPs (*i.e*., not random) could lead to better accuracies, but we leave this as future work.

**Fig. 3:**
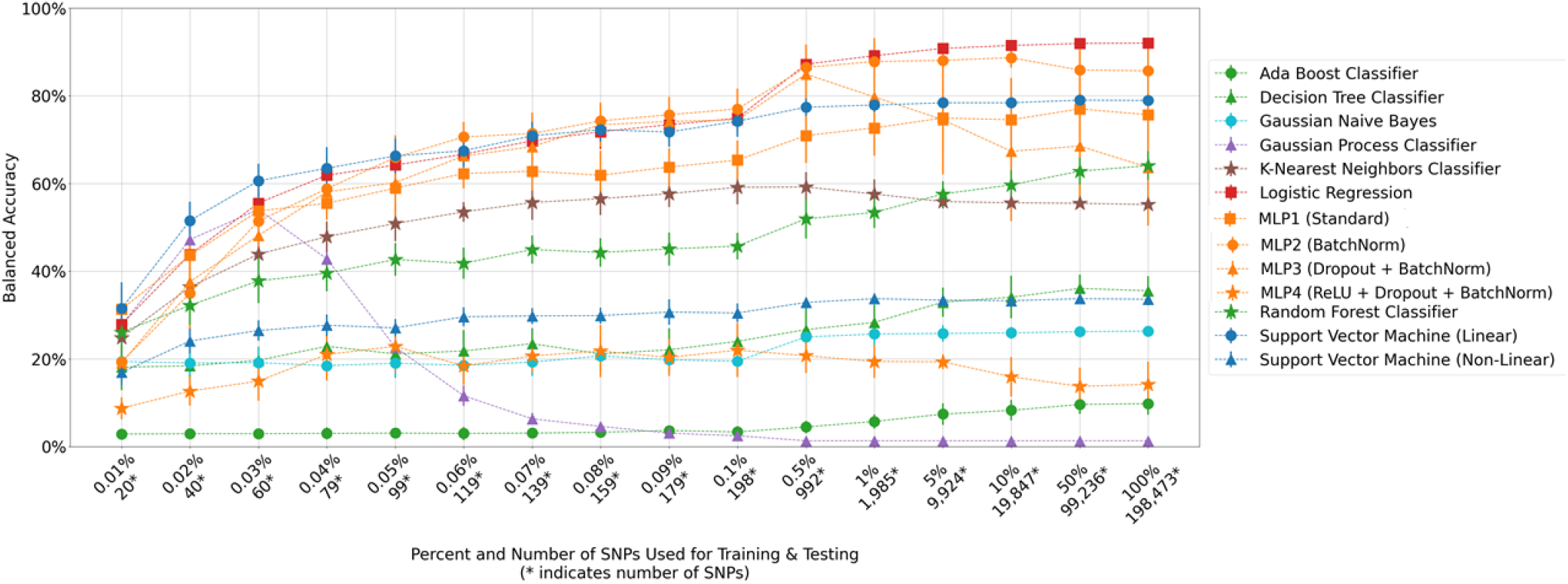
Breed Classification Results. Results plotted in green indicate tree-based and boosting methods; orange indicate Multi-Layer Perceptron (MLP) methods; and blue indicate Support Vector Machine (SVM) methods. All other methods are plotted in different colors.

**Fig. 4:**
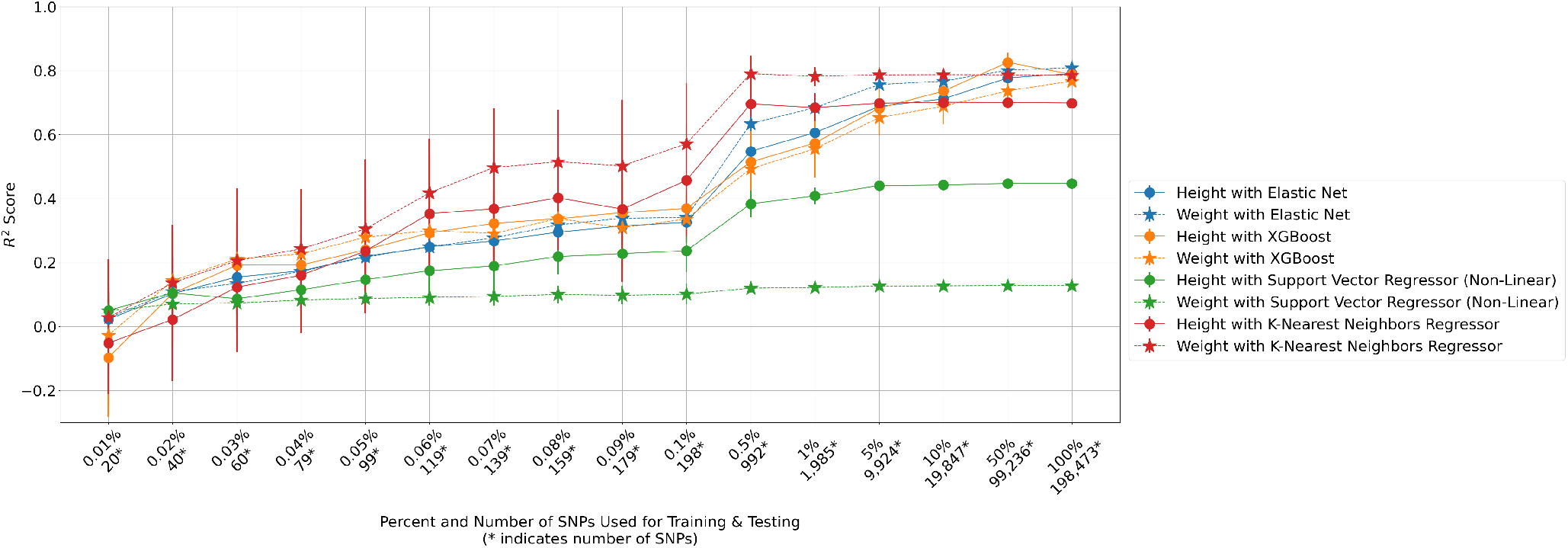
Height and Weight Regression Results. Results for height and weight predictions of a method are shown in the same color.

Figure 3 indicates that many methods achieve success on the breed classification task, and balanced accuracy increases as more SNPs are analyzed. Logistic regression achieves the best performance overall (92% balanced accuracy) when all SNPs are analyzed. The top five methods are all linear: logistic regression, MLP2 (MLP1 with batch normalization), linear SVM, MLP1, and MLP3 (MLP1 with dropout and batch normalization). The non-linear MLP4 is the only MLP network that does not achieve high success on this task. Instead, it performs comparably to other non-linear methods, such as the non-linear SVM and the Decision Tree Classifier, even with the regularization features (*i.e*., batch normalization and dropout) that aid the same architecture in achieving top-5 success. These results are consistent with other work that show linear techniques outperform or match performance of more complex non-linear methods [5], [10], [11]. We believe that linear models outperform non-linear ones because the relationship between dog SNP sequences and breeds, given enough genomic positions, is additive. The linear methods correctly identify dog breeds from genomic sequences, even when only a few SNPs are considered. For example, many methods achieve 50% balanced accuracy after analyzing only 0.5% of SNPs (*i.e*., 992 SNPs). Even when only 0.01% of SNPs (*i.e*., 20 SNPs) are analyzed, linear SVM, MLP1, and logistic regression achieve 31%, 31%, and 28% balanced accuracy, respectively.

In general, balanced accuracy increases as the number of SNPs increases. However, performance of the MLPs decreases slightly when substantially more SNPs are analyzed (*e.g*., 0.5%+ SNPs for MLP3; 50%+ for MLP2). The MLP architecture has a large number of parameters when the inputs are very long sequences and fails to find accurate solutions via gradient descent. Increasing the size of the dataset and using stronger regularization could improve performance for networks for longer sequences. An MLP architecture is highly customizable in terms of its size, regularization techniques, and hyperparameters. Many experiments may be required before a successful MLP architecture and training parameters are found. By comparison, logistic regression requires less design during startup. However, it can suffer from long training times, especially when input sequence lengths are extremely long. Despite the greater challenges in designing a MLP, MLPs are fully differentiable. Thus, they can be combined with other neural networks to create a system with end-to-end backpropogation for other ML tasks. Each method has trade-offs to consider when designing systems for genomic data processing.

Figure 4 shows that most regression methods achieve similar performance for both height and weight prediction and that all methods’ performance increases as more SNPs are analyzed. With only 0.5% of SNPs, KNN achieves an *R*^2^ value of approximately 0.8 and maintains this trend when more SNPs are analyzed. However, for fewer percentages of SNPs, these methods cannot predict phenotypes as well as the classification methods predict breeds. Although the linear methods perform the best for the breed classification task, both non-linear and linear methods achieve comparable success on these regression tasks. In fact, non-linear methods surpass the linear Elastic Net method for most percentages of SNPs. Overall, simple methods (logistic regression and KNN) seem to outperform many of the popular and state-of-the-art techniques (*e.g*., neural networks and boosting trees). Results indicate that phenotype predication can be accomplished with relatively few SNPs for dogs. Considering the full length of dog chromosome data is approximately ∼2.4 billion positions, it is impressive that dog breed can be predicted with 50% balanced accuracy with only 40 randomly selected SNPs. Because dogs have been selectively bred, their genome sequences have been pushed apart. For other organisms that have not been selectively bred, we expect that more SNPs are needed to determine salient information.

## IV. CONCLUSION

We predict dog phenotypes from their genotypes and explore how few SNPs are required to predict a dog’s breed, height, and weight. Because we analyze dog breeds that result from selective breeding, we observe that phenotype prediction can be achieved with as few as 0.5% of SNPs with high accuracy. Even just 0.02% of SNPs achieves 50% balanced accuracy for breed classification. We also observe that linear classification methods outperform non-linear methods, while non-linear methods often match linear methods for regression tasks. We will extend our future work to predict more phenotypes, both of dogs and of other organisms.

